# The major genetic risk factor for severe COVID-19 is inherited from Neandertals

**DOI:** 10.1101/2020.07.03.186296

**Authors:** Hugo Zeberg, Svante Pääbo

## Abstract

A recent genetic association study (Ellinghaus *et al.* 2020) identified a gene cluster on chromosome 3 as a risk locus for respiratory failure in SARS-CoV-2. Recent data comprising 3,199 hospitalized COVID-19 patients and controls reproduce this and find that it is the major genetic risk factor for severe SARS-CoV-2 infection and hospitalization (COVID-19 Host Genetics Initiative). Here, we show that the risk is conferred by a genomic segment of ~50 kb that is inherited from Neandertals and occurs at a frequency of ~30% in south Asia and ~8% in Europe.

## Main text

The coronavirus SARS-CoV-2 pandemic has caused considerable morbidity and mortality, claiming the lives of more than half a million people to date (WHO 2020). The disease caused by the virus, COVID-19, is characterized by a wide spectrum of severity of clinical manifestations, ranging from asymptomatic virus carriers to individuals experiencing rapid progression to respiratory failure (Vetter *et al.* 2020). Early in the pandemic it became clear that advanced age is a major risk factor, as well as male sex and some co-morbidities (Zhou *et al.* 2020). These risk factors, however, do not fully explain why some have no or mild symptoms while others become seriously ill. Thus, genetic risk factors are being investigated. An early study (Ellinghaus *et al*. 2020) identified two genomic regions associated with severe COVID-19: one region on chromosome 3 containing six genes and one region on chromosome 9 that determines the ABO blood group. A recently released dataset from the COVID-19 Host Genetics Initiative finds that the region on chromosome 3 is the only region significantly associated with severe COVID-19 at the genome-wide level (Fig. 1A) while the signal from the region determining ABO-blood group is not replicated (The COVID-19 Host Genetics Initiative 2020).

**Figure 1.**
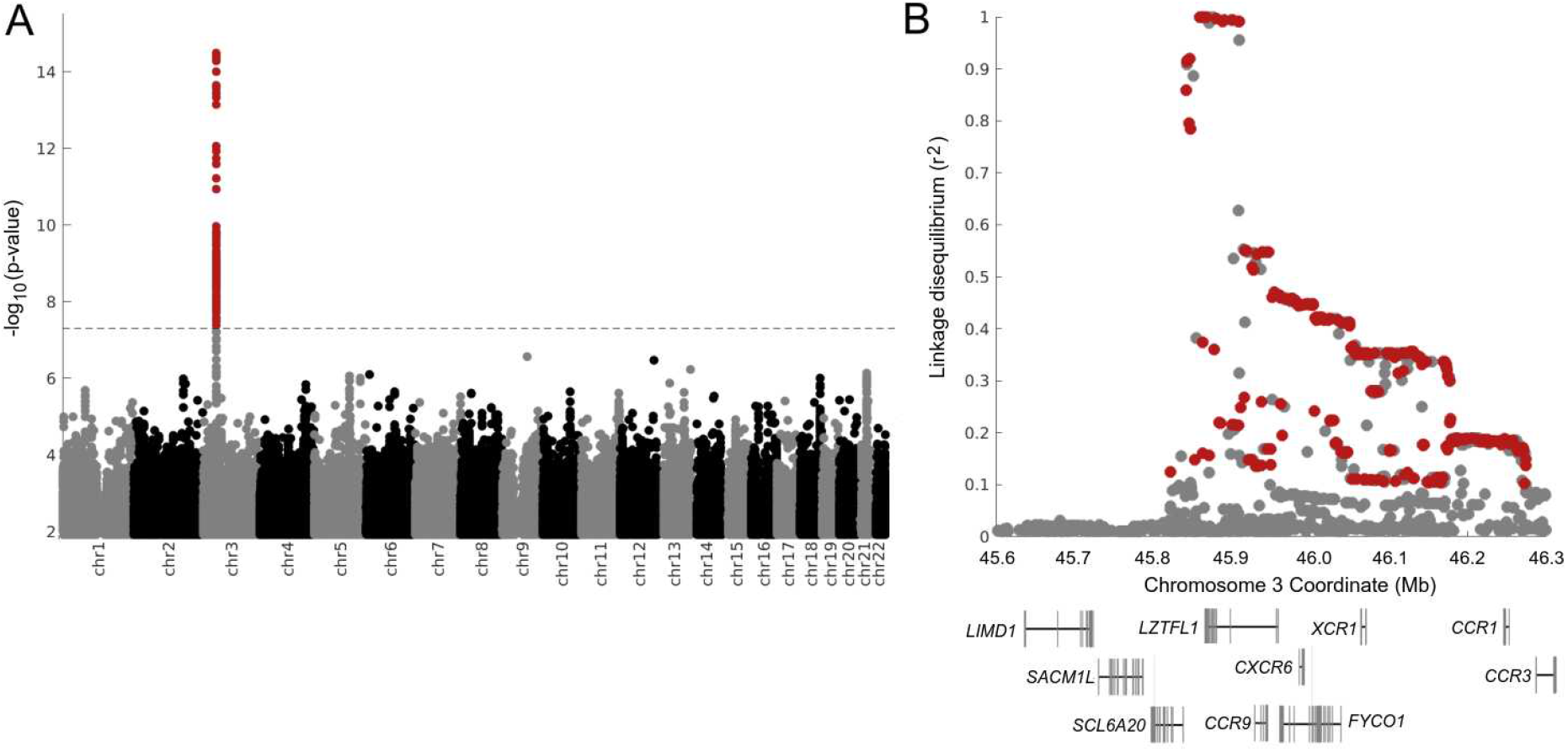
Genetic variants associated with severe COVID-19. **A)** Manhattan plot of a genome-wide association study comprising 3,199 hospitalized COVID-19 patients and 897,488 population controls. Dashed line indicates genome wide significance (p=5e-8). Data and figure modified from the COVID-19 Host Genetics Initiative (https://www.covid19hg.org/) **B)** Linkage disequilibrium between a lead risk variant (rs11385942, Ellinghaus *et al.* 2020) and genetic variants in Eurasian populations. Red marks genetic variants where alleles are correlated to the risk variant (r^2^>0.1) and the correlated alleles match the *Vindija 33.19* Neandertal genome. Note that some individuals carry a Neandertal-like haplotype that extends to ~400 kb. The x axis gives *hg19* coordinates.

The genetic variants which are associated with severe COVID-19 on chromosome 3 (chr3: 45,859,651-45,909,024, *hg19*) are all in high linkage disequilibrium (LD), *i.e.* they are all strongly associated with each other in the population (r^2^>0.99), and span 49.4 thousand bases (kb) (Fig. 1B). A haplotype of such length could be due to positive selection, to an unusually low recombination rate in the region, or to that the haplotype entered the human population by gene flow from Neandertals or Denisovans that occurred some 40,000 to 60,000 years ago (Sankararaman *et al.* 2012). Positive selection seems unlikely at least under current conditions when the odds ratio for respiratory insufficiency upon SARS-CoV-2 for heterozygous carriers of the haplotype is 1.70 (95% CI, 1.27 to 2.26, Ellinghaus *et al.* 2020). The recombination rate in the region is not unusually low (0.53 cM/mb, Kong *et al.* 2002). We therefore investigated whether the haplotype may have come from Neandertals or Denisovans.

The previously identified lead risk insertion variant (rs11385942) (Ellinghaus *et al.* 2020) is present in all 33 DNA fragments covering this this position in the *Vindija 33.19* Neandertal, a ~50,000-old-old Neandertal from Croatia in southern Europe (Prüfer *et al.* 2017). Of 14 single nucleotide variants in the 1000 Genomes Project that are in high LD (r^2^>0.99) with the insertion risk variant in Eurasian populations, 12 occur in a homozygous form in the *Vindija 33.19* Neandertal (Fig. 1B). Four of these variants occur the “Altai” as well as in the *Chagyrskaya 8* Neandertals, both of whom come from the Altai Mountains in southern Siberia and are ~120,000 and ~50,000 years old, respectively (Table S1) while none occur in the Denisovan genome. Thus, the risk haplotype is similar to the corresponding genomic region in the Neandertal from Croatia and less similar to the Neandertals from Siberia.

We next investigated whether the risk haplotype of 49.4 kb might be inherited by both Neandertals and present-day people from the common ancestors of the two groups that lived in the order of 500,000 years ago (Prüfer *et al.* 2014). The longer a present-day haplotype that shared with Neandertals is, the less likely this is to be the case as recombination in each generations will tend to break up haplotypes into smaller segments. Assuming a generational time of 29 years (Langergraber *et al.* 2012), the local recombination rate (0.53 cM/Mb), a split between Neandertals and modern humans of 550,000 years (Prüfer *et al.* 2014), and interbreeding between the two groups ~50,000 years ago, and using a published equation (Huerta-Sánchez *et al.* 2020), we exclude that the Neandertal-like haplotype derives from the common ancestor (p = 0.0009). It thus entered the modern human population from Neandertals. Its close relationship to the Croatian *Vindija 33.19* Neandertal is compatible with that this Neandertal has been shown to be closer to the majority of the Neandertals who contributed DNA to present-day people than the other two Neandertals (Mafessoni *et al*. 2020).

A Neandertal DNA haplotype present in the genomes of people living today is expected to be more similar to a Neandertal genome than to other haplotypes in the current human population. To investigate the relationships of the 49.4 kb-haplotype to Neandertal and to other human haplotypes we analysed all 5,008 genomes in the 1000 Genome Project for this genomic region. We included all positions which are called in the Neandertal genomes and excluded variants found on only one chromosome and haplotypes seen only once in the 1000 Genomes data. This resulted in 253 present-day haplotypes containing 450 variable positions. Figure 2 shows a phylogenetic tree relating such haplotypes found more than 10 times (see Fig. S1 for all haplotypes). We find that all risk haplotypes associated with the risk for severe COVID-19 form a clade with the three high-coverage Neandertal genomes. Within this clade, they are most closely related to the *Vindija 33.19* Neandertal.

**Figure 2.**
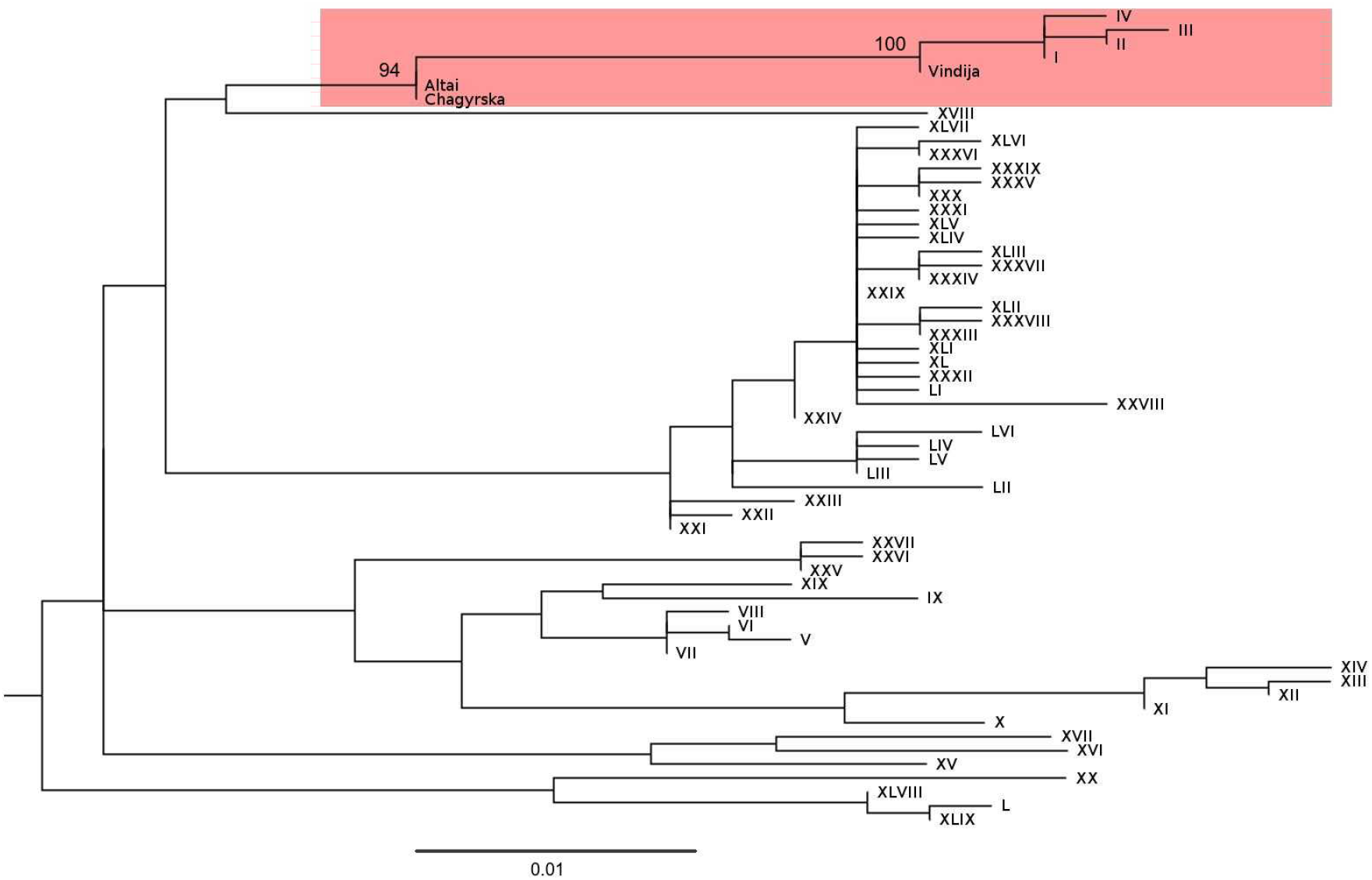
Phylogenetic tree relating DNA sequences covering the core Neandertal haplotype in 1000G individuals. The coloured box indicates Neandertal and risk haplotypes for severe COVID-19. Arabic numbers indicate bootstrap support (100 replicates). Phylogenies were constructed using maximum-likelihood and the Hasegawa-Kishino-Yano-85 model (Hasegawa *et al.* 1985). The tree is rooted with the inferred ancestral sequence of present-day humans from Ensembl (Yates *et al.* 2020). There are no heterozygous positions in this region in the three Neandertal genomes.

**Figure 3.**
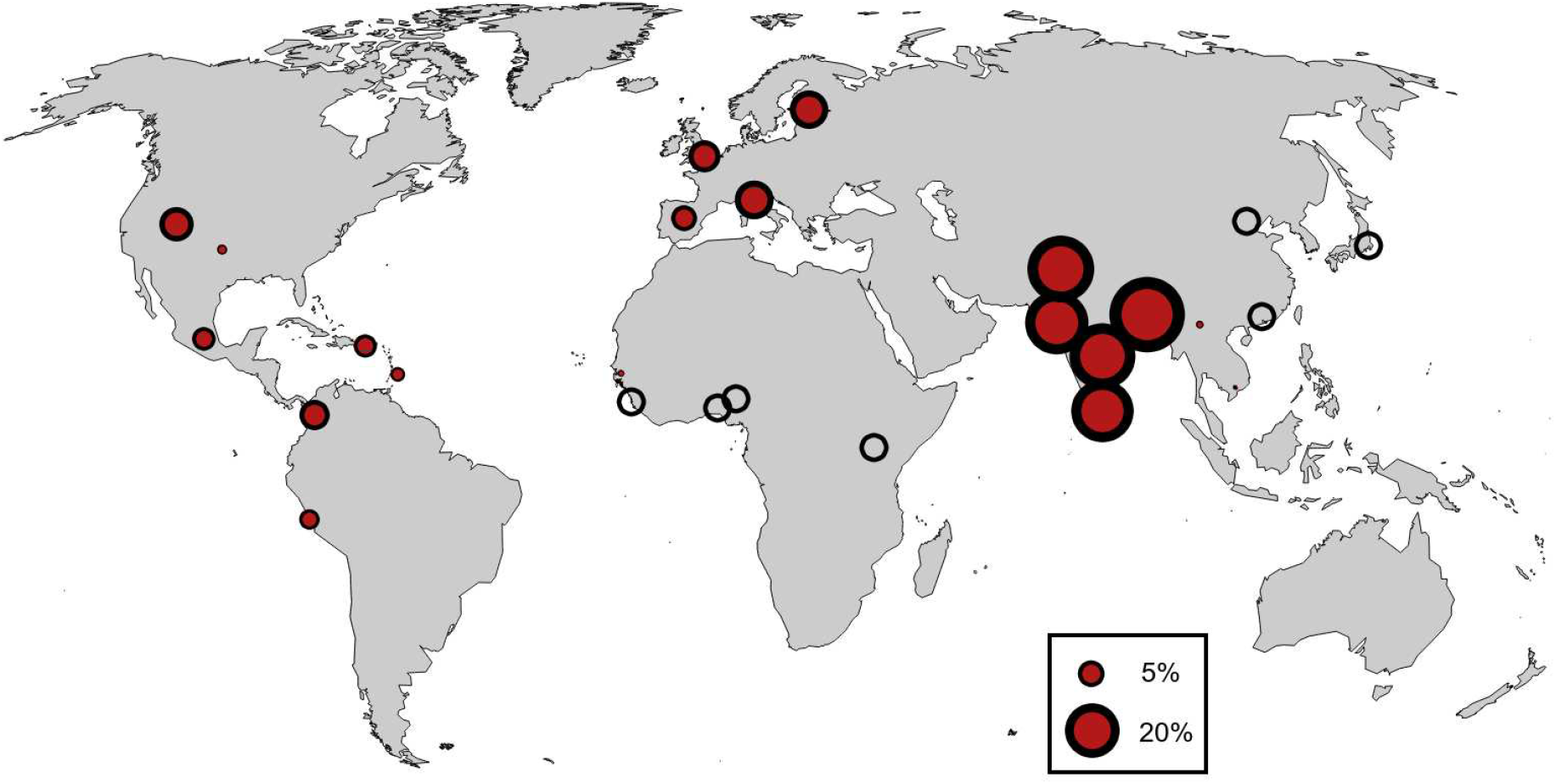
Geographical distribution of the Neandertal core haplotype conferring risk for severe COVID-19. Empty circles denote populations where the Neandertal haplotype is missing. Data from the 1000 Genomes Project.

Among the individuals in the 1000 Genome Project, the Neanderthal-derived risk haplotypes is almost completely absent in Africa, consistent with that gene flow from Neandertals into African populations was limited and probably indirect (Chen *et al.* 2020). The Neandertal haplotype occurs in South Asia at a frequency of 30%, in Europa at 8%, among admixed Americans at 4% and at lower frequencies in East Asia. The highest frequency occurs in Bangladesh, where more than half the population (63%) carries at least one copy of the Neandertal risk variant and 13% is homozygous for the variant. The Neandertal variant may thus be a substantial contributor to COVID-19 risk in certain populations.

Currently it is not known what feature in the Neandertal-derived region confers risk for severe COVID-19 and if the effects of any such feature is specific to current coronaviruses or indeed to any other pathogens. Once this is elucidated, it may be possible to speculate about the susceptibility of Neandertals to relevant pathogens. However, in the current pandemic, it is clear that gene flow from Neandertals has tragic consequences.

## Acknowledgements

We are indebted to the COVID-19 Host Genetics Initiative (HGI) for making the GWAS data available, to the Max Planck Society and the NOMIS Foundation for funding, and to Nordforsk for funding to sequence patients for the COVID-19 HGI.

## Supplementary material

**Figure S1.**
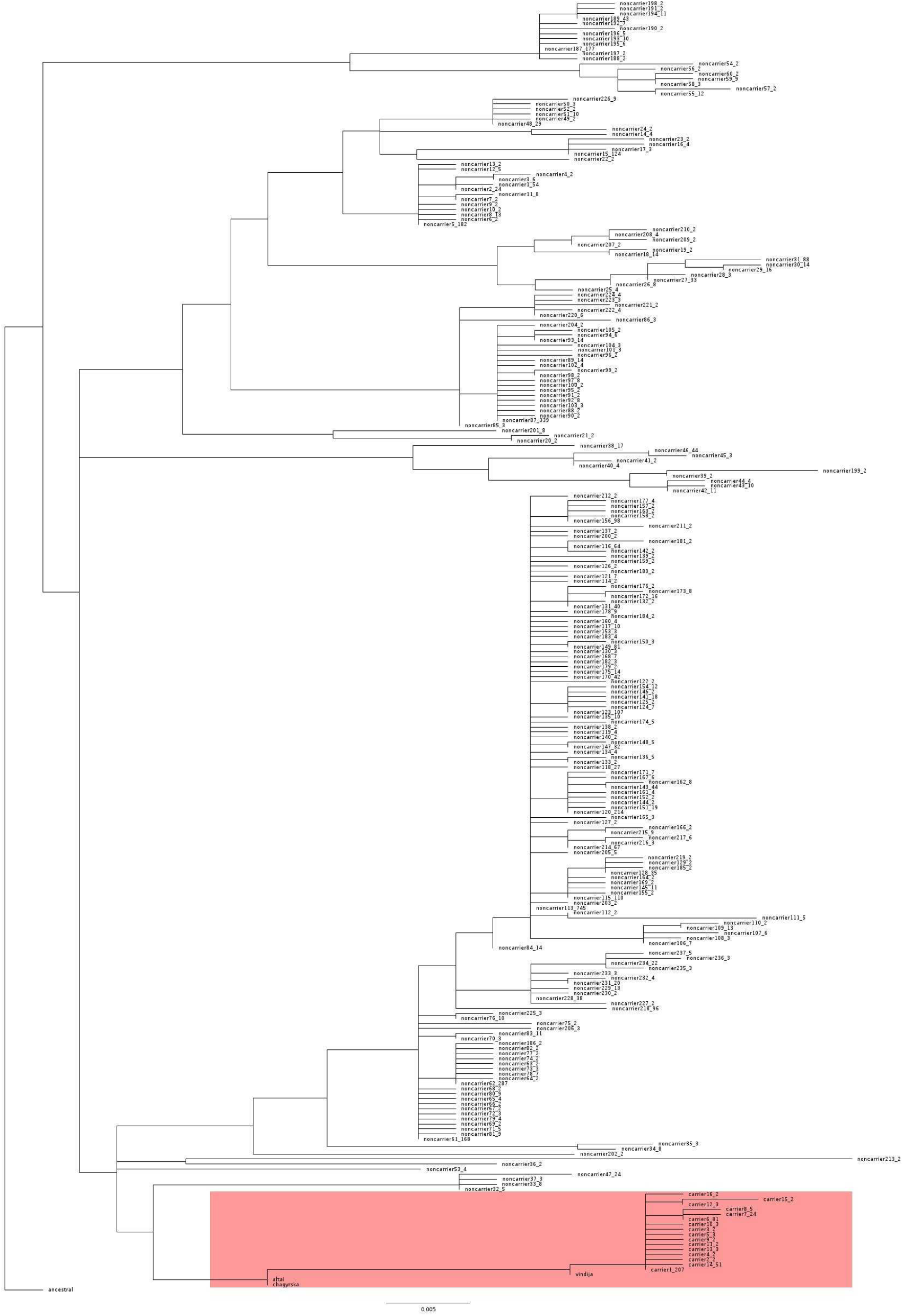
A phylogenetic tree estimating the relationships of all haplotypes in 1000 Genomes individuals covering the genomic region of the core Neandertal risk haplotype. Haplotypes marked as carriers contain the minor allele of rs10490770, which is in perfect LD in Eurasian populations with the leading risk polymorphisms. All carriers fall in a clade with the Neandertal genomes. The last numbers in the haplotype designations indicate the numbers of chromosomes carrying the haplotype.

**Table S1.**
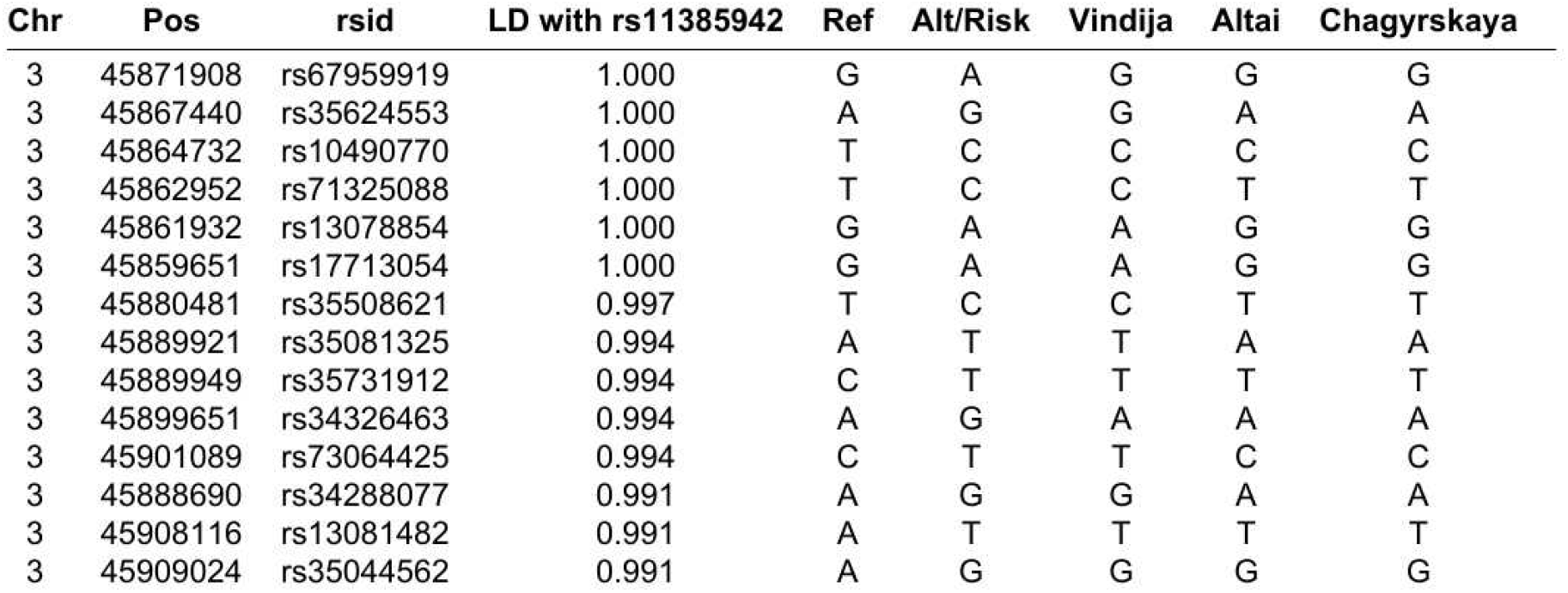
Genetic variants in LD (r^2^>0.99) with rs11385942 in Eurasian populations and the corresponding Neandertal variants. Data from the 1000 Genomes Project. “Ref” gives the *hg19* alleles.

